# Mineral crystal thickness in calcified cartilage and subchondral bone in healthy and osteoarthritic knees

**DOI:** 10.1101/2021.06.15.448181

**Authors:** Mikko A.J. Finnilä, Shuvashis Das Gupta, Mikael J. Turunen, Iida Kestilä, Aleksandra Turkiewicz, Viviane Lutz-Bueno, Elin Folkesson, Mirko Holler, Neserin Ali, Velocity Hughes, Hanna Isaksson, Jon Tjörnstrand, Patrik Önnerfjord, Manuel Guizar-Sicairos, Simo Saarakkala, Martin Englund

**Author notes:** Shared senior authors (Simo Saarakkala, Martin Englund).

## Abstract

Osteoarthritis (OA) is the most common joint disease globally. In OA, articular cartilage degradation is often accompanied with sclerosis of the subchondral bone. However, the association between OA and tissue mineralization at the nanostructural level is currently not understood. Especially, it is technically challenging to identify calcified cartilage, where relevant but poorly understood pathological processes like tidemark multiplication and advancement occur. Here, we used state-of-the-art micro-focus small-angle X-ray scattering with high 5µm spatial resolution to determine mineral crystal thickness in human subchondral bone and calcified cartilage. Specimens with a wide spectrum of OA severities were acquired from the medial and lateral compartments of medial compartment knee OA patients (*n*=15) and cadaver knees (*n*=10). For the first time, we identified a well-defined layer of calcified cartilage associated with pathological tidemark multiplication, containing 0.32nm thicker crystals compared to the rest of calcified cartilage. In addition, we found 0.2nm thicker mineral crystals in both tissues of the lateral compartment in OA compared with healthy knees, indicating a loading-related disease process since the lateral compartment is typically less loaded in medial compartment knee OA. Furthermore, the crystal thickness of the subchondral bone was lower with increasing histopathological OA severity. In summary, we report novel changes in mineral crystal thickness during OA. Our data suggest that unloading in the knee is associated with the growth of mineral crystals, which is especially evident in the calcified cartilage. In the subchondral bone, mineral crystals become thinner with increasing OA severity, which indicates new bone formation with sclerosis.

**One Sentence Summary:** Mineral crystal thickness increases with osteoarthritis in the lateral condyle that is typically unloaded.

## INTRODUCTION

In large joints, rigid bone provides a structural foundation for smooth articular cartilage that enables almost frictionless movement. The most important constituents of articular cartilage are proteoglycans and collagens, mainly of type II. Charged proteoglycans attract ions, which generate osmotic pressure and thus, have a crucial role in maintaining the dynamic compressive properties of cartilage*(1, 2)*. Fibrillar collagen is organized in a pattern also known as ‘Benninghoff arcades’ and it provides tensile strength for the cartilage *(3)*. These properties allow cartilage to dissipate and distribute loading, protecting the underlying subchondral bone*(4, 5)*. The subchondral bone is composed of type I collagen fibrils embedded with hydroxyapatite crystals, contributing to the toughness and stiffness of the tissue*(6–8)*. The subchondral bone should be rigid enough to transfer mechanical load but elastic enough to reduce high stress in articular cartilage*(9)*. Furthermore, bone is a metabolically active tissue that has the capability to adapt to mechanical loading, and it has been identified as a viable target tissue for pharmaceutical therapies*(10)*.

Calcified cartilage forms an interface between the subchondral bone and articular cartilage. A thin basophilic line in histological sections is known as tidemark, which forms the interface between the articular cartilage and calcified cartilage*(11)*. There is another highly mineralized interface between calcified cartilage and subchondral bone called the cement line*(12)*. Calcified cartilage resembles both tissues by consisting of a cartilage-like organic matrix with proteoglycans embedded in a type II collagen network, which is mineralized as the bone tissue*(13, 14)*. Although calcified cartilage has been reported to have a higher degree of mineralization than subchondral bone*(15)*, the hardness and stiffness of the tissue are between those of subchondral bone and articular cartilage*(16, 17)*.

Osteoarthritis (OA) is a degenerative disease that targets most joint tissues. Osteochondral tissues have hierarchical structures that change over multiple length scales. In articular cartilage, early disease processes involve chondrocyte apoptosis/proliferation and surface fibrillation/edema, which develops to surface discontinuity accompanied by proteoglycan loss. In moderate OA, vertical fissures appear in the articular cartilage followed by cartilage matrix erosion with more evident proteoglycan loss and possibly some collagen formation*(18)*. Subchondral bone sclerosis is a known hallmark of OA. We have suggested that the thickening of bone structures is a synchronous process with cartilage degeneration during OA progression*(19)*.

The calcified cartilage undergoes structural modifications during OA. Perhaps the most evident of the changes is described as the tidemark multiplication*(20, 21)*, which has been linked to the advancement of calcified cartilage towards the articular cartilage*(21–23)*. The uppermost tidemark defines the calcification front presenting tidemark multiplication, which increases the thickness of calcified cartilage. However, two processes could also make calcified cartilage thinner. First, a process similar to endochondral ossification, where deep calcified cartilage is replaced by bone*(24, 25)*, increases the subchondral bone thickness, and thins the calcified cartilage*(26)*. Second, erosion continues on the mineralized surfaces once all cartilage has been removed with denudation with severe OA*(18)*. Considering all of the structure modifying processes it is logical that the thickest calcified cartilage has been observed with moderate OA*(19, 27, 28)*.

Micro-structural changes in bone and calcified cartilage contribute to sclerosis (*stiffening*) of the structures beneath articular cartilage. Furthermore, the mineralization level modulates the stiffness of subchondral tissues*(29)*. Mineral crystal thickness is known to change with tissue maturation*(30)* and is also linked to mechanical properties*(31)*. However, the nano-structural changes in bone and especially in calcified cartilage that are associated with OA are much less clear. Therefore, we aimed to quantify the mineral crystal thickness employing state-of-the-art micro-focus small-angle X-ray scattering (μSAXS) in calcified cartilage and subchondral bone, comparing end-stage knees of OA patients to healthy knees of deceased donors (**Fig. 1**). We hypothesized that mineral crystals are thicker in calcified cartilage than subchondral bone and that mineral crystal thickness increases with OA progression and age in both calcified cartilage and subchondral bone.

**Fig. 1.**
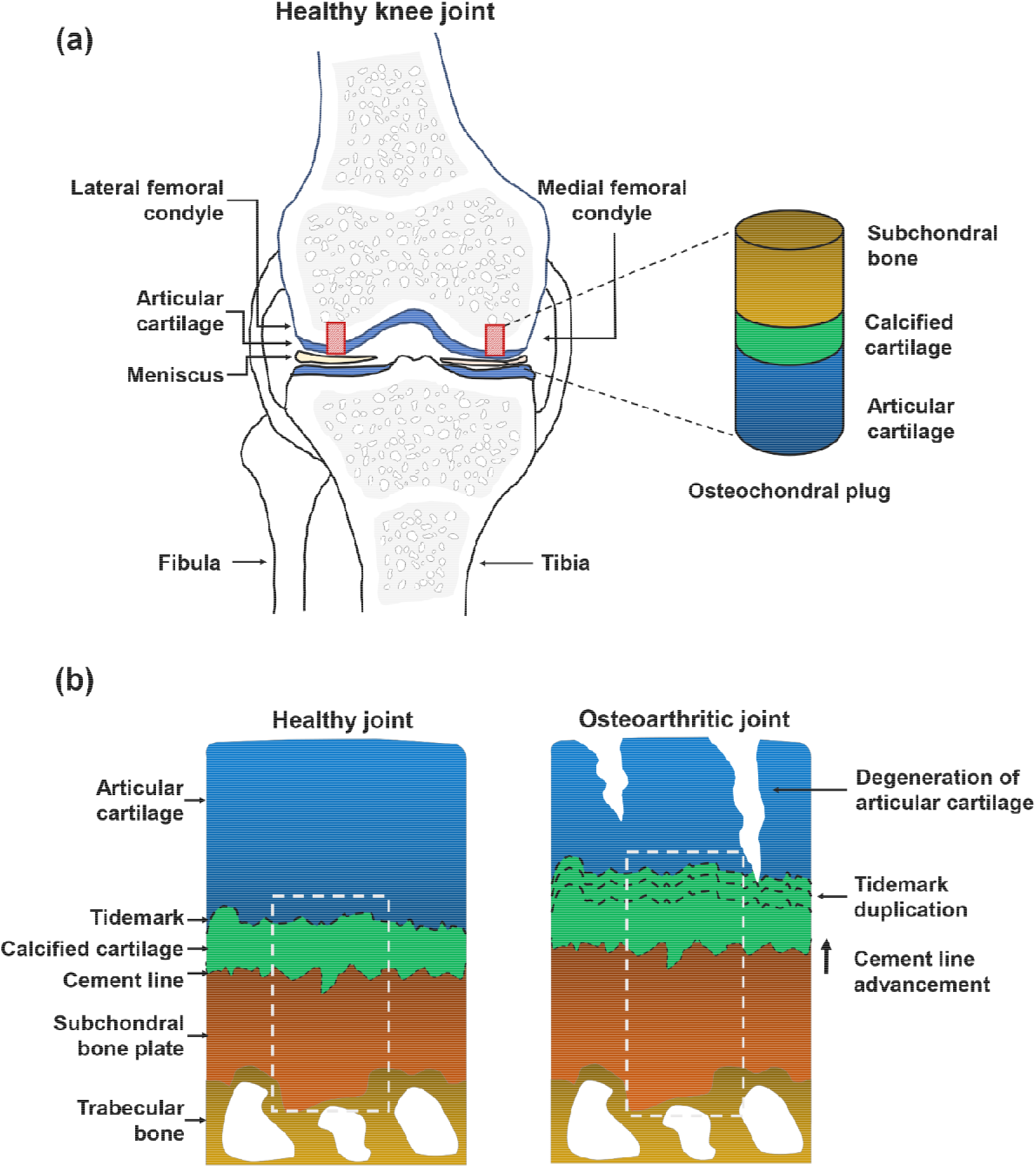
Schematic representation of osteochondral sample collection and μSAXS measurement of healthy and osteoarthritic joints. (A) Osteochondral plugs were collected from both lateral and medial femoral condyles. (B) Schematic cross-sectional representation of samples taken from healthy and osteoarthritic joints. The white dotted line represents the 500 μm wide μSAXS measurement area extending from the deep articular cartilage to the subchondral bone (500-1000 μm).

## RESULTS

### Sample donor descriptive

We collected samples from fifteen patients, who underwent total knee replacement (TKR), due to advanced OA in the medial compartment. We compared these to samples from ten donors with healthy joints (**Table S1**). Our study design contained four groups of osteochondral samples from load-bearing areas of medial and lateral femoral condyles from TKR patients and deceased donors that will henceforth be referred to as i) Medial^TKR^ (*n*=15), ii) Lateral^TKR^ (*n*=15), iii) Medial^donor^ (*n*=10), and iv) Lateral^donor^ (*n*=10). The age range of donors was greater than TKR patients. The heights of patients and donors were similar in both sexes.

### Histopathological severity of OA and tidemark multiplication

We performed a histopathological evaluation of osteochondral samples according to the OARSI grading system*(18)*. The histopathological evaluation solely focuses on changes of the articular cartilage from mild to advanced OA, and subchondral bone modifications are only accounted for in the final grades. Since most of the histopathological evaluation accounts for only articular cartilage modifications, it can be considered as the golden standard for cartilage degeneration. In the cadaveric donors, we observed relatively healthy cartilage with intact surface or focal fibrillation through the superficial zone (**Fig. 2A**). Five donor samples showed features of mild OA and there was a presence of vertical fissures or even cartilage matrix loss. All Medial^TKR^ samples had a loss of cartilage matrix and in some cases, there were also erosions of calcified cartilage. The OARSI grade distribution in the lateral compartment of TKR patients resembled the grade distribution in the medial side of the cadaveric donors.

**Fig. 2.**
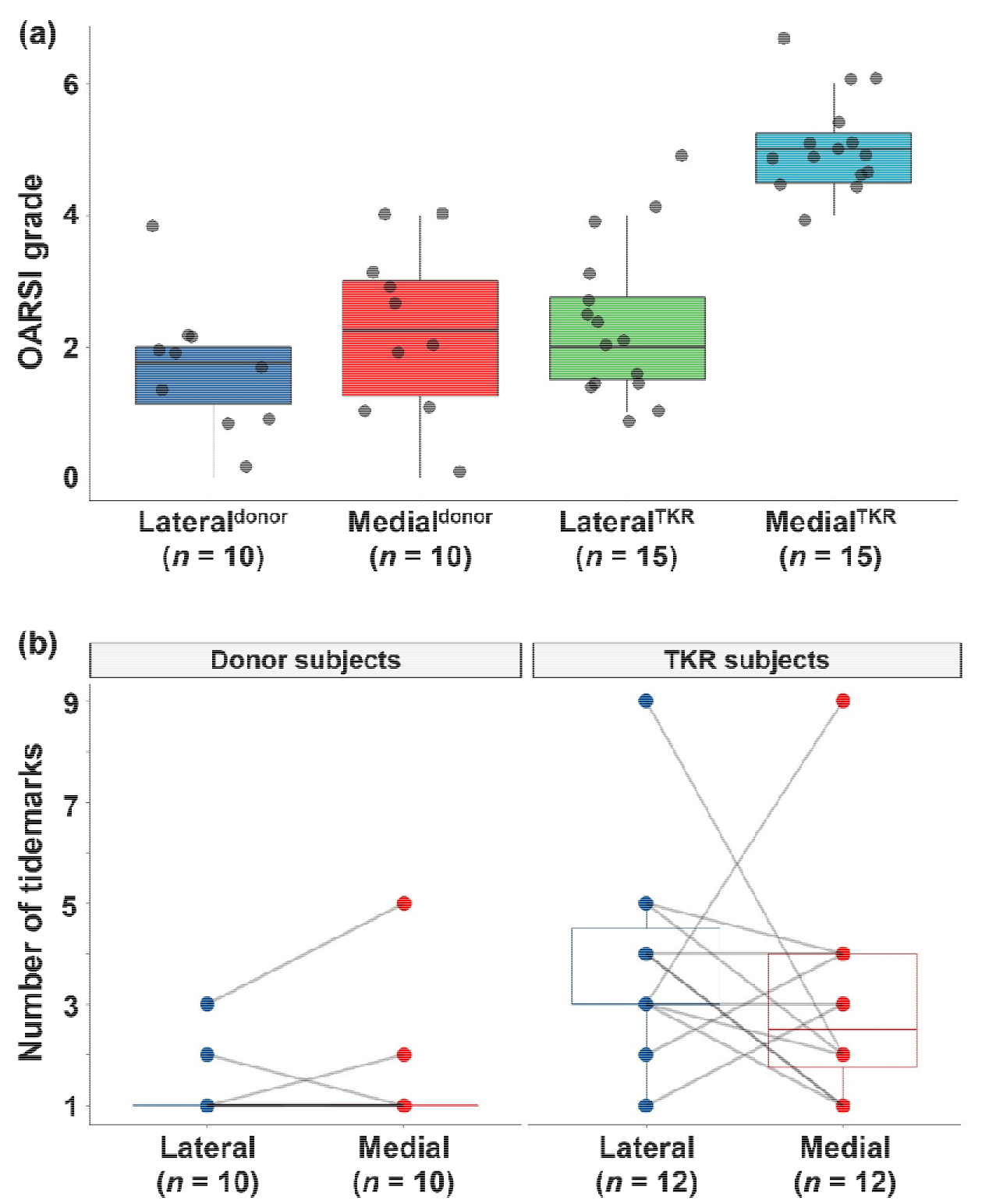
Histopathology of the osteochondral samples. (A) Boxplot with jitter showing the OARSI grades (with sub-grades) in the lateral and medial condyles of cadaveric donors and total knee replacement (TKR) patients. Each grade represents the following key feature. Grade 0: fully intact cartilage; Grade 1: Intact surface with cellular changes and/or edema; Grade 2: Surface discontinuity; Grade 3: Vertical fissures; Grade 4: cartilage erosion; Grade 5: denudation (articular cartilage matrix loss to calcified cartilage); and Grade 6: deformation. (B) Boxplot with a pairwise comparison showing the number of tidemarks, which were counted from the histopathological images. There was complete erosion of calcified cartilage in three samples from the Medial^TKR^ group and these patients were excluded from the pairwise comparison.

In addition to OARSI grading, we counted the number of tidemarks from histopathological images. The number of tidemarks reflects the tidemark advancement, a process in which calcified cartilage extends into articular cartilage*(18)*. We observed a greater number of tidemarks in TKR patients compared to the donors. However, we could not identify any clear pattern for tidemark multiplication between the medial and lateral compartments (**Fig. 2B**).

### Mineral crystal thicknesses in the osteochondral junction

To visualize mineral crystal thickness distribution in mineralized tissues we utilized the state-of-the-art cSAXS beamline at the Swiss Light Source. We collected scattering patterns with 5 μm spatial resolution. The mineral crystal thickness was then analyzed by performing a weighted iterative curve fitting on I(*q*) scattering derived by azimuthally integrating recorded scattering patterns.

Calcified cartilage and subchondral bone were segmented by utilizing unsupervised *K*-means clustering (**Fig. 3**), based on the fitted I(*q*) curves. This allowed spatially resolving of mineral crystal thicknesses separately for calcified cartilage and subchondral bone (**Fig. S1 and Fig. S2**). Segmentation was verified by visually comparing various cSAXS visualizations with histological images. Furthermore, we observed that the upper-most cluster(s) in calcified cartilage was co-localized with a layer between two top-most tidemarks in the calcified cartilage (**Fig. S3)**. Interestingly, the mineral crystals in the top-most layer appear to be thicker compared to the rest of the calcified cartilage (**Table 1** and **Fig. 4**). We propose that this layer represents the active mineralization front during tidemark advancement.

**Fig.3.**
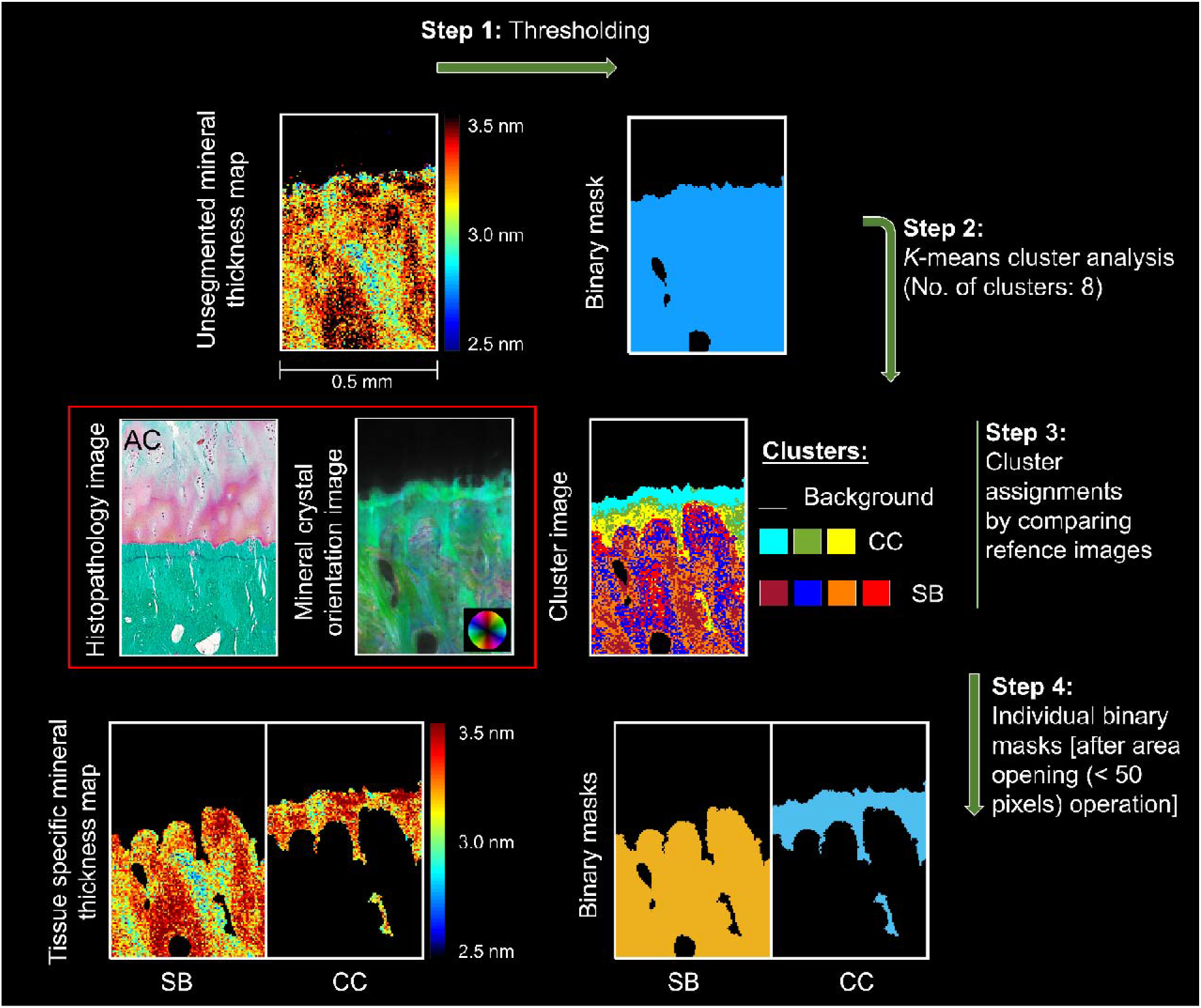
Tissue segmentation from 2D μSAXS images. Step 1: the mineral crystal thickness maps were binarized using an absolute threshold on scattering intensity. Step 2: *K*-means cluster analysis of the μSAXS data (I(*q*) scattering curves at each measurement point). Step 3: Eight clusters were found optimal for most of the samples when comparing the cluster image to the crystal orientation image. The cluster assignment was further confirmed by comparing the cluster image to the histopathological images. Step 4: final tissue-specific binary masks for calcified cartilage (CC) and subchondral bone (SB) were generated after de-speckling.

**Table 1.** Descriptive statistics of the mineral crystal thickness of two layers in the calcified cartilage. Results are displayed as means (standard deviations) of the mineral crystal thicknesses of the upper-most mineralization front and the rest of calcified cartilage along with the difference (with 95% confidence interval) between them, in both medial and lateral compartments, respectively.

| Compartment location | Mineral crystal thickness (nm) |  | Difference (95% CI) |
| --- | --- | --- | --- |
|  | Upper-most mineralization front of CC | Rest of CC |  |
| **Lateral (*n* = 15)** | 3.64 (0.29) | 3.27 (0.17) | 0.36 (0.26, 0.47) |
| **Medial (*n* = 8)** | 3.36 (0.28) | 3.13 (0.09) | 0.23 (0.01, 0.46) |

**Fig. 4.**
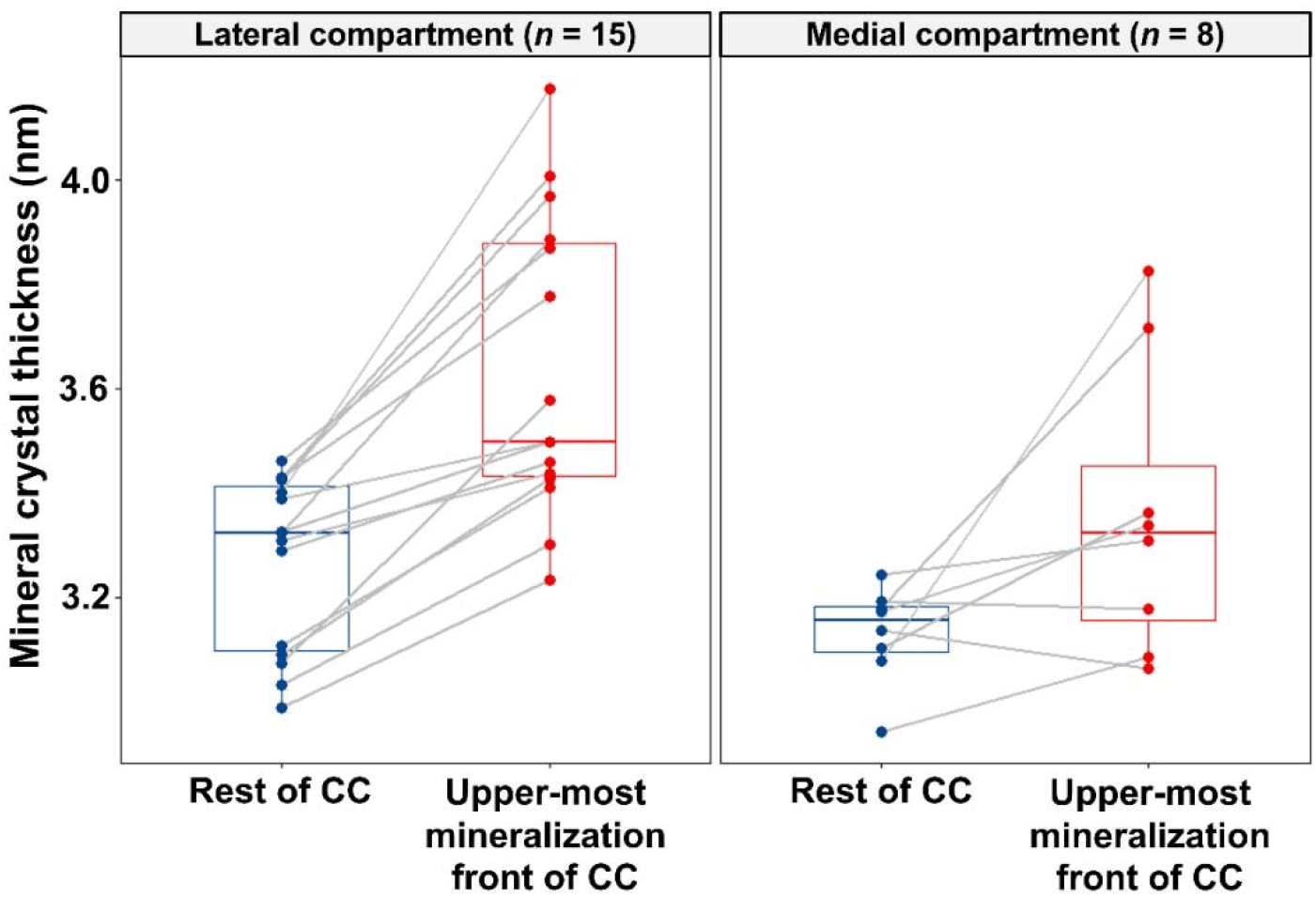
Mineral crystal thickness in the two layers of calcified cartilage (CC). Boxplot showing the pair-wise comparison of the mineral crystal thickness between the upper-most mineralization front in the CC and the rest of the CC, from both lateral and medial compartments, in the osteochondral specimens with multiple tidemarks (fifteen and eight specimens for lateral and medial compartment, respectively).

### Mineral crystal thicknesses in cadaveric donors and total knee replacement patients

Mineral crystal thickness in the subchondral bone was around 3.0 nm in all other groups except the lateral compartment of the TKR patients with mineral crystal thickness of 3.19 nm (**Table 2**). In all groups, the mineral crystal thicknesses in calcified cartilage are higher compared to those in the subchondral bone.

**Table 2.** Descriptive statistics of the tissue-specific mineral crystal thickness. Results are displayed as means (standard deviations) of the mineral crystal thicknesses of the calcified cartilage and the subchondral bone together with the difference (with 95% confidence interval) between them, in both medial and lateral compartments of cadaveric donors (donor) and total knee replacement (TKR) patients.

| Group | Mineral crystal thickness (nm) |  | Difference (95% CI) |
| --- | --- | --- | --- |
|  | Calcified cartilage | Subchondral bone |  |
| **Lateral<sup>donor</sup>** | 3.15 (0.18) | 3.01 (0.22) | 0.14 (0.05, 0.23) |
| **Medial<sup>donor</sup>** | 3.19 (0.21) | 2.96 (0.16) | 0.23 (0.10, 0.35) |
| **Lateral<sup>TKR</sup>** | 3.41 (0.16) | 3.19 (0.19) | 0.22 (0.11, 0.33) |
| **Medial<sup>TKR</sup>** | 3.24 (0.10) | 3.03 (0.12) | 0.21 (0.14, 0.29) |

In the subchondral bone, the difference between Lateral^TKR^ and Medial^TKR^ was 0.17 nm (95%CI 0.03, 0.31), and between Lateral^donor^ and Lateral^TKR^ 0.17 nm (95%CI 0.08, 0.26). However, this difference diminishes after adjusting for age (**Fig. 5**). The mineral crystal thickness in calcified cartilage was the highest in the Lateral^TKR^ and the difference after adjusting for age and BMI remained at 0.2 nm (95%CI 0.10, 0.29) and 0.23 nm (95%CI 0.11, 0.35) when comparing to Medial^TKR^ or Lateral^donor^, respectively (**Fig. 5**).

**Fig. 5.**
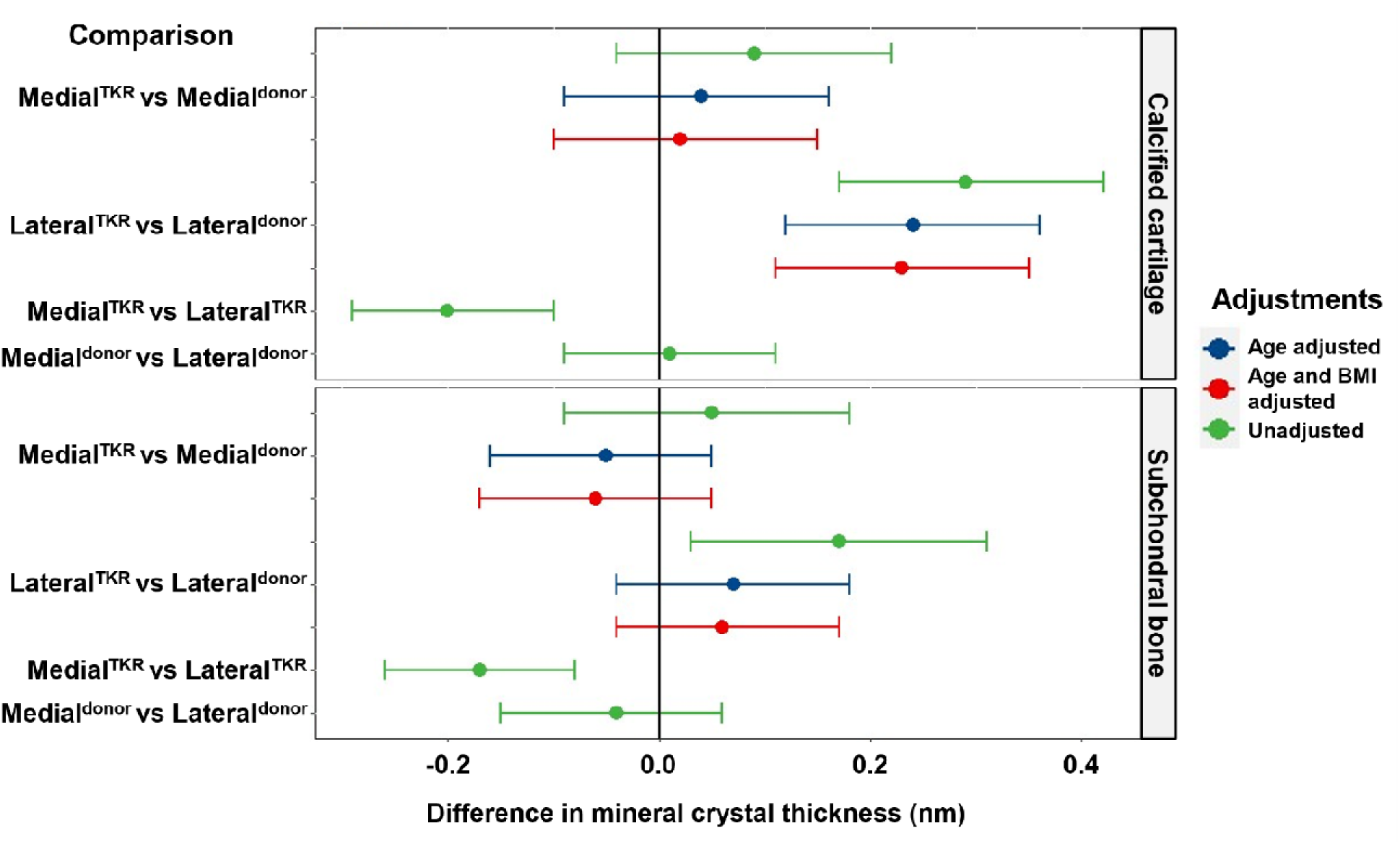
Compartment-specific comparison of the mineral crystal thickness between cadaveric donors (donor) and total knee replacement (TKR) patients. Differences in mineral crystal thickness displayed with a 95% confidence interval in the calcified cartilage and the subchondral bone. The model was adjusted for age, and then for age and BMI. The comparison between medial and lateral compartments from the same knee is adjusted for all person-and knee-level confounding through the design and use of a mixed-effects model.

To evaluate possible changes in mineral crystal thickness with histopathological OA severity, we performed an association analysis between OARSI grade and mineral crystal thickness. After adjusting for compartment location, patient status, age, and BMI, we found a decrease of −0.04 nm (95%CI −0.07, −0.02) per unit increase in OARSI grade but only in the subchondral bone (**Fig. 6**). The slope in calcified cartilage was −0.01 (95%CI −0.05, 0.03) suggesting no relevant association.

**Fig. 6.**
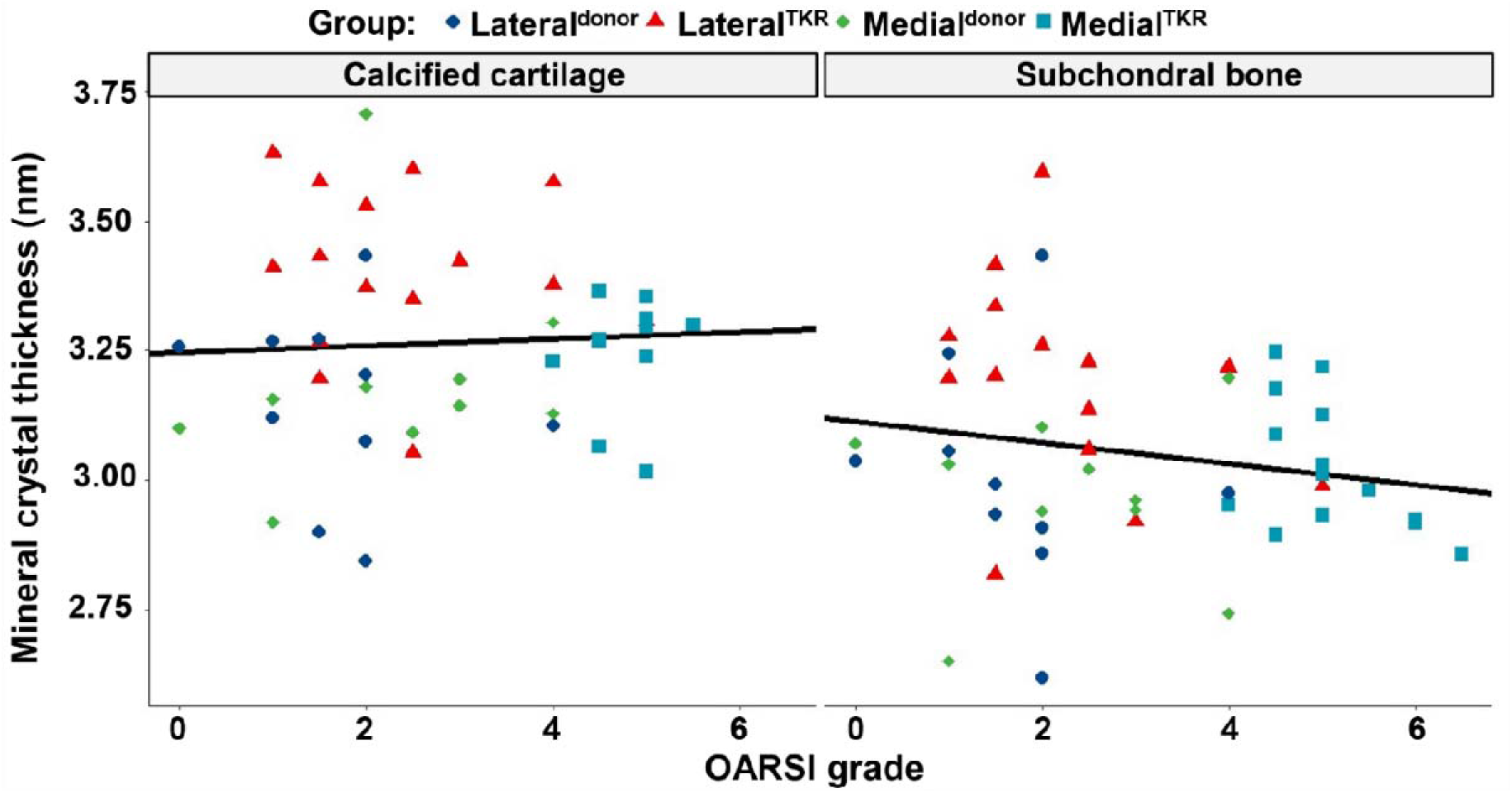
Associations between the mineral crystal thickness and histopathological osteoarthritis progression. Scatter plots with the regression lines (unadjusted) showing the associations between the mineral crystal thickness and the OARSI grade, in the calcified cartilage and the subchondral bone, respectively. Association was found only for the subchondral bone with a reduction of 0.04 nm for each OARSI grade.

## DISCUSSION

In this study, we spatially mapped the mineral crystal thickness across the osteochondral junction using state-of-the-art micro-focus small-angle X-ray scattering (μSAXS). Results were aligned with our first hypothesis that the mineral crystals are thicker in calcified cartilage than in subchondral bone. Against our second hypothesis, we observed a reduction in the thicknesses of mineral crystals with osteoarthritis severity but only in the subchondral bone. However, for the first time, we observed an increase in mineral crystal thickness of both osteochondral tissues in the lateral compartment of total knee replacement patients compared to donors, where cartilage shows less degeneration compared to the medial side.

The geometry, as well as orientation and architecture of the mineral crystals in osteochondral mineralized tissues, are all indicative of the strength of the tissues*(32)*. Early studies on the ultrastructure of bone using electron microscopy in the 1950s showed that the mineral crystals are plate-shaped and roughly aligned with their long axes parallel to the long axes of the collagen fibrils*(33, 34)*. Besides plate-like geometry, needle-like geometry has also been proposed in several studies*(35, 36)*. Modern evidence using a combination of 3D electron tomography and 2D electron microscopy suggests that a fractal-like hierarchical bone architecture, where needle-shaped mineral crystals merge laterally to form platelets that organize into stacks of roughly parallel platelets (thickness 5 nm) bridging multiple collagen units*(37)*.

The thickness of the mineral crystal is commonly assumed to be regulated by the structure of the organic matrix*(38)* and by non-collagenous proteins*(39)*, and the fact that the calcified cartilage and subchondral bone have different organic matrices, it is crucial to study the mineral crystal thickness of calcified cartilage and subchondral bone, individually, as a function of OA and age. However, owing to limitations in the spatial resolution or contrast of current nondestructive imaging modalities, possible changes in the mineral crystal thickness of calcified cartilage and subchondral bone due to OA have not been studied separately until now. Zizak *et al.* reported that the calcified cartilage possesses similar mineral crystal thickness to that of underlying bone by using SAXS analysis*(40)*. They proposed that the mineral crystal thickness of these subchondral tissues might be governed by the non-collagenous proteins. In contrast, here, we report greater mineral crystal thickness in calcified cartilage compared to subchondral bone. This finding is aligned with our previous investigation of the human osteochondral junction by using Raman microspectroscopy, where a higher degree of mineralization and more stoichiometric perfect crystal lattice was found in the calcified cartilage compared to the subchondral bone plate*(15)*. Accordingly, we propose that the thicker mineral crystal might be related to the fewer non-stoichiometric substitutions of carbonate for phosphate ions in the crystal lattice of calcified cartilage compared to the bone. The carbonate substitution in the apatite crystal lattice is reported to decrease mineral crystallinity, as well as cause contraction of the a-axis and expansion of the c-axis dimensions of the unit cell*(41)*. Furthermore, the other dimensions *i.e.,* length and width, and the volume of crystals typically increase with increased crystal thickness*(42)*. This could indicate that larger crystals are either more tightly packed or stronger Raman scatterers.

Fawns and Landell in 1953 were among the first to describe ‘tidemark’ as the most recent calcified border of calcified cartilage, using histological staining*(43)*. The tidemark has a wavy morphology, which helps to reduce stress concentrations on the calcified cartilage. The presence of two or more tidemarks (tidemark multiplication) is a sign of advancement of calcified cartilage into articular cartilage during OA progression and which may be related to the subchondral resorption activity*(44)*. There appears to be a similar endochondral ossification process in the calcified cartilage that occurs in long bones during growth*(45, 46)*. In this ossification process, chondrocytes near the tidemark are shown to have a hypertrophic phenotype, express type X collagen*(47)*, and become mineralized. Poorly crystalline calcium inorganic orthophosphate nanocrystals are deposited around the chondrocytes in the organic matrix of the cartilage promoting the advancement of calcified cartilage*(48)*. The non-continuous but periodic process of upward mineralization of articular cartilage, indicated by tidemark multiplication was demonstrated in the femoral heads of mature rabbits already in 1971 by Lemperg*(49)* and in osteoarthritic human joints by Green *et al.(50)*. In the present study, there were more tidemarks in TKR patients compared to donors. Furthermore, previous studies have reported a higher level of mineralization near tidemark than in the region of calcified cartilage near the bone with quantitative backscattered electron imaging*(12, 51)*. When we compared our cluster image from the μSAXS analysis with respective histological images of adjacent sections, we related the upper-most cluster to the layer of calcified cartilage separated by two top-most tidemarks. We propose that this layer is indicative of the most recently mineralized layer of calcified cartilage due to ossification. Remarkably, we observed the thickest mineral crystal in this topmost calcified cartilage (**Fig. 4**). This could not be observed in our previous Raman study*(15)* and to our knowledge, this has not been reported before in the literature. This may indicate that during the tidemark advancement the mineral crystals have greater thickness after deposition, and they re-structure to thinner plates with maturation. Eventually, the calcified cartilage substance is removed by osteoclasts (carried by capillaries) and then replaced by bone substance synthesized by osteoblasts during new bone formation*(48)*.

The present study allows a detailed spatial comparison of mineral crystal thickness and OA severity assessed with histopathology. We compared both lateral and medial compartments of the femoral osteochondral junction from TKR patients with end-stage medial compartment knee OA and deceased donors without known OA. We found that the lateral compartment of the TKR patients has thicker mineral crystals in both subchondral bone and calcified cartilage, compared to the OA-affected medial compartment, as well as the lateral compartment of donors (**Fig. 5**). Importantly, these changes occur in the osteochondral mineralized tissues of the lateral compartment while the greatest cartilage degeneration occurs in the medial compartment. Thus, the difference between TKR patients and donors found in mineral crystal thickness in the lateral compartment’s calcified cartilage indicates that mineralization is an independent process compared to cartilage degeneration. This is further supported by the fact that mineral crystal thickness is higher in calcified cartilage in the Lateral^TKR^ compared to Lateral^donor^ even after adjusting for age and BMI. In subchondral bone, age adjustment reduces the difference in group-wise comparison. Finally, we observed an association between OARSI grade and mineral crystal thickness in subchondral bone, suggesting that both age and histopathological OA progression affect mineral crystal thickness in the bone. The latter would support our previous study where the microstructural bone changes are associated with cartilage degeneration*(19)*. Thus, it appears that the subchondral bone undergoes structural changes over various hierarchical levels with OA progression.

Owing to the fact that the TKR patients had end-stage knee OA on the medial compartment, we expected to find changes in mineral crystal thickness in this compartment. Surprisingly, we observed increased mineral crystal thickness in the lateral compartment, with apparently “healthy” cartilage. In individuals with a healthy knee joint, during the gait cycle in normal walking, the load distribution is not equal between the medial and lateral tibiofemoral compartments and the medial compartment exhibits higher loads*(52, 53)*. The medial shift of the load-bearing axis due to varus alignment further increases loading across the medial compartment*(53, 54)*. It has been reported that the absolute load (normalized to body weight) is increased in OA and due to mediolateral loading distribution, a relatively high proportion of the load is subjected over the medial compartment*(55)*. In the same study, half of the subjects with medial compartment knee OA demonstrated unloading of the lateral compartment in mid to late stance*(55)*. An increase in joint loading and cartilage stress are commonly considered to be related to bone architecture alteration and cartilage degeneration, respectively*(46)*. Previous studies showed that abnormal loads may start a remodeling response in bone*(56, 57)* and induce changes in composition and mechanical properties in cartilage*(58)*, respectively. Thus, the increased mineral crystal thickness in the lateral compartment could be related to biomechanics. This is further supported by a study from O’Connor who reported increased mineral apposition and tidemark advancement after experimental unweighting of hind limbs in rats*(59)*.

We have previously proposed that mineralization is related to the remodeling of subchondral bone*(15)*. The bone remodeling and modeling of the subchondral bone are integral parts of OA progression*(23, 28, 60)*. The remodeling process involves the interplay between the osteoclasts and osteoblasts and the bone tissue itself would become younger with the increased remodeling rate. The newer bone tissue typically has lower mineralization*(28, 29)* and mineral crystal thickness*(30, 61, 62)*. In this study, the mineral crystal thickness in subchondral bone was lower in knees with higher histopathological OA severity (**Fig. 6**). Such an increase in subchondral bone remodeling could be related to abnormal joint loading*(9)* or osteoclastic activation under the hormonal influence*(63)*. Osteoclasts may generate vascular channels through calcified cartilage*(44, 59)*. Studies on surgically induced animal models of OA suggest that the reduction in subchondral plate thickness in early OA is associated with increased bone remodeling*(26, 64)*. New (primary) bone formation beneath the intact articular cartilage during OA progression was observed using a bovine patella model system*(22)*, which suggests the bone formation at the osteochondral junction might be a possible indicator of early OA and change the permeability of this crucial interface. Although in the early OA, bone resorption predominates in the process of bone remodeling, but in the late OA, bone resorption decreases without reduced bone formation*(28)*. Thus, the thickening of the subchondral plate along with subchondral sclerosis in the late OA stages are observed*(19, 27, 28, 65, 66)*.

In summary, we observed increased mineral crystal thickness in calcified cartilage and subchondral bone of the unloaded compartment in the patients with the medial compartment knee OA. These results indicate that the mineralization of osteochondral tissues is an independent process from cartilage erosion, and it could be driven by a lack of local mechanical loading. Our findings indicate that loading is an important factor for biomineralization of calcified cartilage during OA development, and could be exploited when developing tissue engineering strategies for the osteochondral interface.

## MATERIALS AND METHODS

### Tissue sample preparation

This study was approved by the regional ethical review board (Lund University; Dnr 2015/39 and Dnr 2016/865). From the MENIX biobank, osteochondral samples from load-bearing areas of medial and lateral femoral condyles were selected from 15 patients with end-stage medial compartment knee OA (8 women, 7 men) who had a TKR at Trelleborg Hospital, Sweden (**Table S1**). Obtained femoral condyles were frozen at −80°C within 2h of extraction and delivered on dry ice to the biobank in Lund for further storage at −80°C. During the TKR surgery, the surgeon’s Outerbridge classification*(67)* of the knee joint cartilage was required to be a grade IV in the medial compartment and grade 0 or I in the lateral compartment for the patient to be classified to have medial compartment OA.

In addition, we selected osteochondral samples from load-bearing areas of medial and lateral femoral condyles from 10 deceased human donors (5 women, 5 men), also from the MENIX biobank, with donors obtained from Skåne University Hospital, Lund, Sweden. No known diagnosis of knee OA or rheumatoid arthritis was allowed for the donors, who were included in this study. The femoral condyles were obtained within 48 hours post-mortem and frozen at – 80°C within 2h of extraction.

On the sample preparation day, we thawed medial and lateral femoral condyles from both TKR and donor samples which were trimmed in sample diameter to 5mm with a trephine drill or low-speed diamond saw, respectively. We fixed the specimens in 4% saline-buffered formaldehyde for 7 days at 4° C. Subsequently, we fixed the samples in 70% ethanol for 24h. Specimen processing continued with dehydration in ascending ethanol concentrations (80% – 95% – 100% – 100%), each step lasting for at least 6h. After dehydration, the specimens were embedded in polymethylmethacrylate. To deactivate the monomer, we filtered Methyl Methacrylate (MMA) through a column filled with aluminum oxide. We infiltrated samples first with MMA in a solution containing deactivated MMA, 1.4 (vol%) Nonyphenyl polyethylenenglycol acetate, and 5.5μg/ml Benzoyl peroxide at 4°C for 24h, after which the solution was replaced with a fresh one. In the final step, the container was filled with the previous solution with an additional 5μl/ml of N,N,Dimethyl-p-toluidine for plastification that was completed in the following 24h at 4°C.

### Histological analyses

Six histological sections (3-μm-thick) were cut from the PMMA blocks followed by Safranin-O or Goldner trichrome staining. The stained sections were used for the histopathological OARSI grading*(18)* and counting the number of tidemarks, respectively. Grading was first conducted independently by two readers [MAJF, IK; inter-observer reliability: ICC (95% confidence interval (CI)) 0.85 (0.69, 0.93) for medial osteochondral samples and 0.79 (0.58, 0.90) for lateral osteochondral samples], and their consensus grade was given as a final grade for each sample.

### Mineral crystal thickness mapping using small-angle X-ray scattering

The μSAXS measurements were performed at the cSAXS – X12SA: Coherent Small-Angle X-ray Scattering beamline at the Swiss Light Source, Paul Scherrer Institute (Villigen, Switzerland). We cut 5 μm thick sections adjacent to the histological sections and fixed those on Kapton tape. Photon energy was set to 12.4 keV and the beam at the sample position was focused to 5×5 μm^2^. SAXS patterns were recorded with Pilatus 2M detector*(68)* placed 7.1 m from the sample and using 30 ms exposure time per point. A 500 μm wide area extending from the mineralized front (tidemark) to the bone marrow (500-1000 μm) was scanned with a 5 μm step size in a continuous line-scan mode. Recorded scattering patterns were azimuthally integrated to obtain the intensity *I* as a function of scattering vector 0, I(*q*) scattering curves *(69)*. The mineral crystal thickness at each point was evaluated from the I(*q*) curves, using weighted iterative curve fittings in the *q*-range of 0.32–1.40 nm^−1^ *(42, 70)*. In the model, the dimensions of mineral crystal are assumed such that the scattering takes place from the mineral plates with finite thickness and infinite size in two other dimensions. Details of the model are explained elsewhere*(42, 70)*. Calcified cartilage and subchondral bone were identified by using unsupervised *K*-means clustering (**Fig. 3**) of the fitted I(*q*) curves. We verified cluster results by comparing adjacent histological sections and mineral crystal orientation images*(69)*. Typically, eight clusters provided meaningful segmentation, but if the reference images indicated incorrect tissue identification, then the number of clusters was iteratively increased until accurate identification was achieved or a maximum number of 20 clusters was reached.

### Statistical analyses

The distribution of mineral crystal thickness was close to normal for almost all samples. Thus, the mean of the distribution was used for further statistical evaluation. The mineral crystal thickness between groups was compared by using a linear mixed-effects model with the patient/donor as a random effect. The patient status (donor vs TKR), the compartment (medial vs lateral), and their interactions were included in the model. Additionally, we adjusted the model for age and BMI. Due to the small sample size, Kenward and Roger’s method was used for estimating the number of degrees of freedom. Moreover, similar linear mixed-effects models were used to estimate the differences in the mineral crystal thickness between the tissue types (calcified cartilage and subchondral bone), as well as between two different layers of calcified cartilage (upper-most mineralization front and rest of calcified cartilage). When calculating the difference between the tissue types, we set both sample groups, tissue types, and their interactions as fixed effects and patient/donor as a random effect. Furthermore, we set both compartment locations, calcified cartilage layers, and their interactions as fixed effects and patient/donor as a random effect to estimate the difference between two different layers of calcified cartilage. We present the estimates with 95% CI. To estimate the association between OARSI grade and mineral crystal thickness, we used a mixed linear regression model adjusted for compartment, patient status, age, and BMI.

## Supplementary Materials

Table S1. Main descriptive of OA patients and donors.

Fig. S1. Segmented mineral crystal thickness maps of the osteochondral junction from the medial femoral condyles.

Fig. S2. Segmented mineral crystal thickness maps of the osteochondral junction from the Lateral femoral condyles.

Fig. S3. Segmentation of the upper-most mineralization front of the calcified cartilage (CC) in samples with multiple tidemarks.

## Acknowledgments

We acknowledge the Paul Scherrer Institut, Villigen PSI, Switzerland for the provision of synchrotron radiation beamtime at the cSAXS beamline X12SA of the SLS. Moreover, we thank Ms. Tarja Huhta for preparing the histological sections.

## Funding

Finnish Cultural Foundation (North Ostrobothnia Regional Fund No. 60172246) (MAJF)

Finnish Foundation for Technology Promotion (IK)

European Union’s Horizon 2020 research and innovation program under the Marie Skłodowska-Curie grant agreement no. 713645 (SDG)

Academy of Finland (Grants No. 268378 and 303786) (SS)

European Research Council under the European Union’s Seventh Framework Program (FP/2007-2013; ERC Grant Agreement No. 336267) (SS)

Swedish Research Council (ME)

The Swedish Rheumatology Association (ME)

Österlund Foundation (ME)

Governmental funding of clinical research within the national health services (ME)

## Author contributions

Conceptualization: MAJF, MJT, SS, ME

Methodology: JT, MAJF, EF, NA, VH, PÖ

Software: MAJF, MJT, SDG, AT

Investigation: MAJF, IK, MJT, VL, MH, MG, SDG, HI, AT

Visualization: SDG

Supervision: SS, ME

Writing – original draft: MAJF, SDG

Writing – review & editing: all

## Competing interests

Authors declare that they have no competing interests.

## Data and materials availability

Data used for this study can be made available by submitting on reasonable request to the corresponding author.

## Notes

### Competing Interest Statement

The authors have declared no competing interest.

