## Supplementary material for "Mineral crystal thickness in calcified cartilage and subchondral bone in healthy and osteoarthritic knees": Supplementary.docx

**Table S1. Main descriptive of OA patients and donors.**

|  | | Patients | | | | Donors | | |
| --- | --- | --- | --- | --- | --- | --- | --- | --- |
|  |  | Total | | Female | Male | Total | Female | Male |
|  |  | **15** | | **8** | **7** | **10** | **5** | **5** |
| Age (years) | Median (range) | 65  (50-79) | | 61  (50-70) | 71  (61-79) | 55  (18-77) | 61  (18-77) | 52  (43-70) |
| Height (cm) | Median (range) | 170  (156-186) | 165  (156-174) | | 177  (170-186) | 174  (155-190) | 162  (155-178) | 177  (173-190) |
| Weight (kg) | Median (range) | 85  (63-121) | | 86.5  (63-103) | 85  (75-121) | 72  (52-127) | 56  (52-67) | 107  (77-127) |
| BMI (kg/m^2^) | Median (range) | 29  (22-37) | | 32.5  (22-37) | 29  (26-35) | 25.5  (16-32) | 23  (16-26) | 33  (25-42) |

**Fig. S1.** **Segmented mineral crystal thickness maps of the osteochondral junction from the medial femoral condyles.** Segmented maps of the mineral crystal thickness from the calcified cartilage (CC) and the subchondral bone (SB) from both cadaveric donors and total knee replacement (TKR) patients are presented, respectively. The respective OARSI grades of the samples are added on top of each map. Width: W_1_, W_2_, and W_3_ are equal to 500 µm.

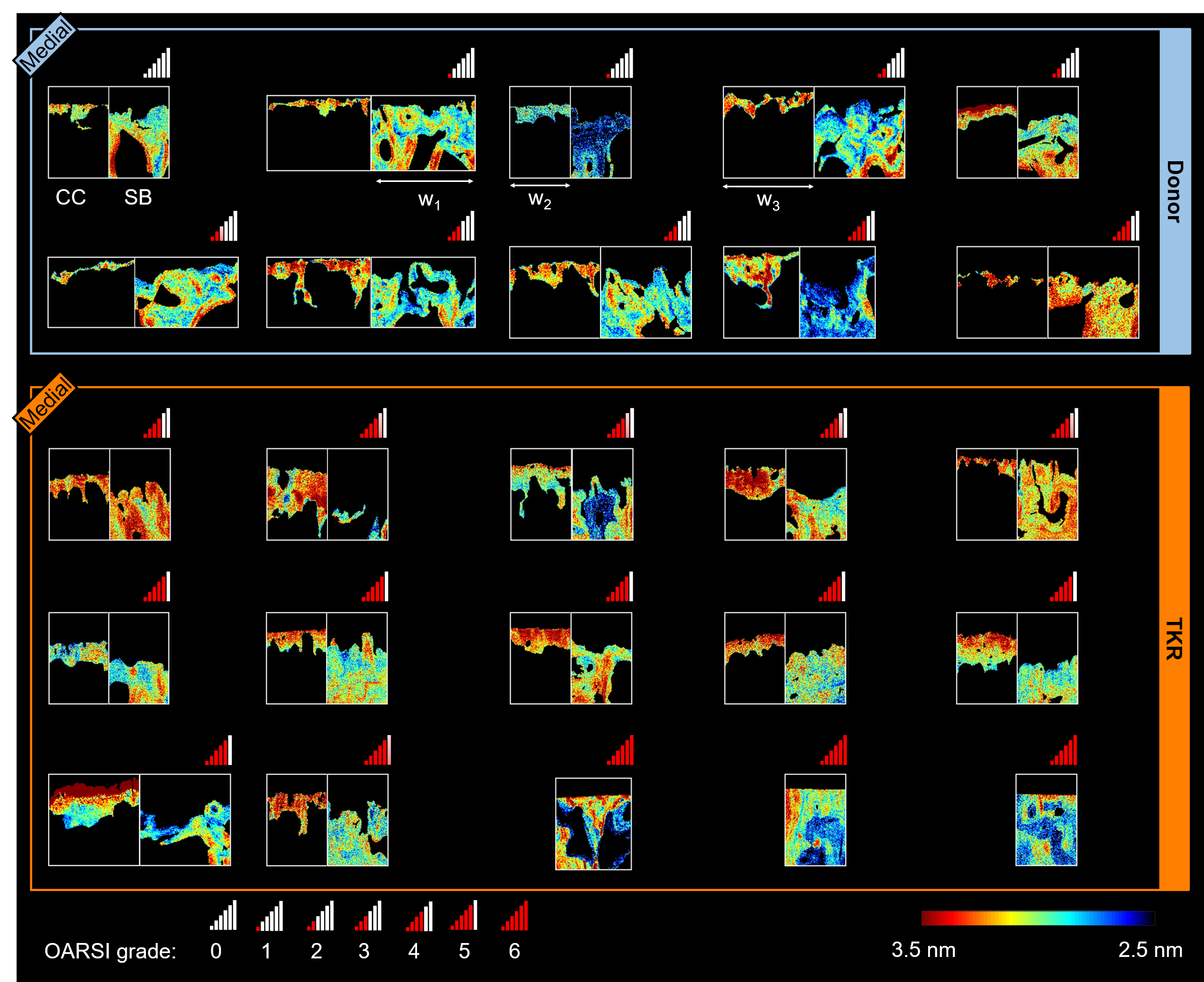

**Fig. S2.** **Segmented mineral crystal thickness maps of the osteochondral junction from the Lateral femoral condyles.** Segmented maps of the mineral crystal thickness from the calcified cartilage (CC) and the subchondral bone (SB) from both cadaveric donors and total knee replacement (TKR) patients are presented, respectively. The respective OARSI grades of the samples are added on top of each map. Width: W_1_ and W_2_ are equal to 500 µm.

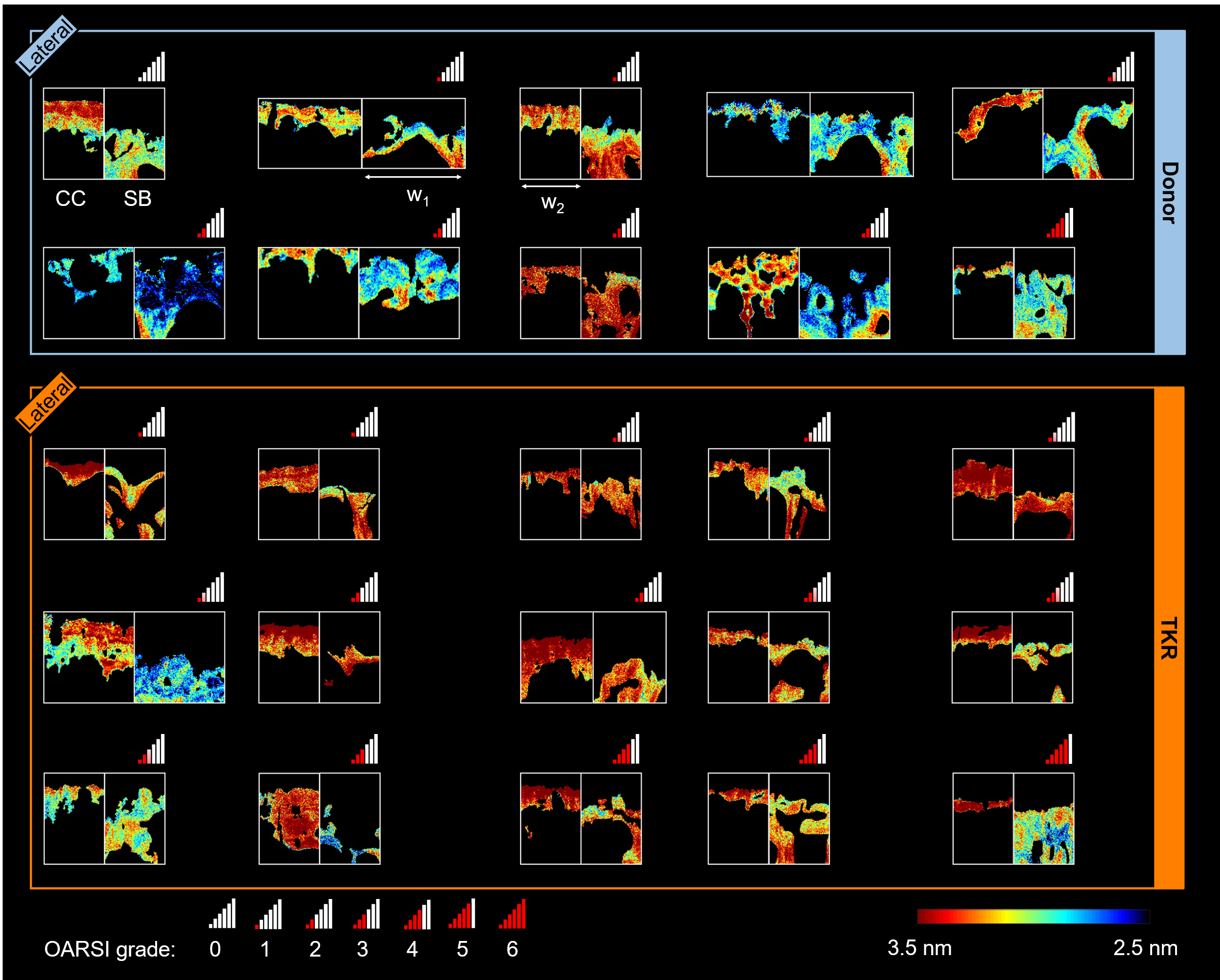

**Fig. S3.** **Segmentation of** **the upper-most mineralization front of the calcified cartilage (CC) in samples with multiple tidemarks.** By comparing the *K*-means cluster image with the mineral crystal orientation image and the adjacent histopathological image, the upper-most cluster of the CC was assigned as the top mineralization front, which is also the closest to the deep articular cartilage (AC). Binary masks of the upper-most mineralization front and rest of CC were generated after despeckling. Abbreviation: SB = subchondral bone.

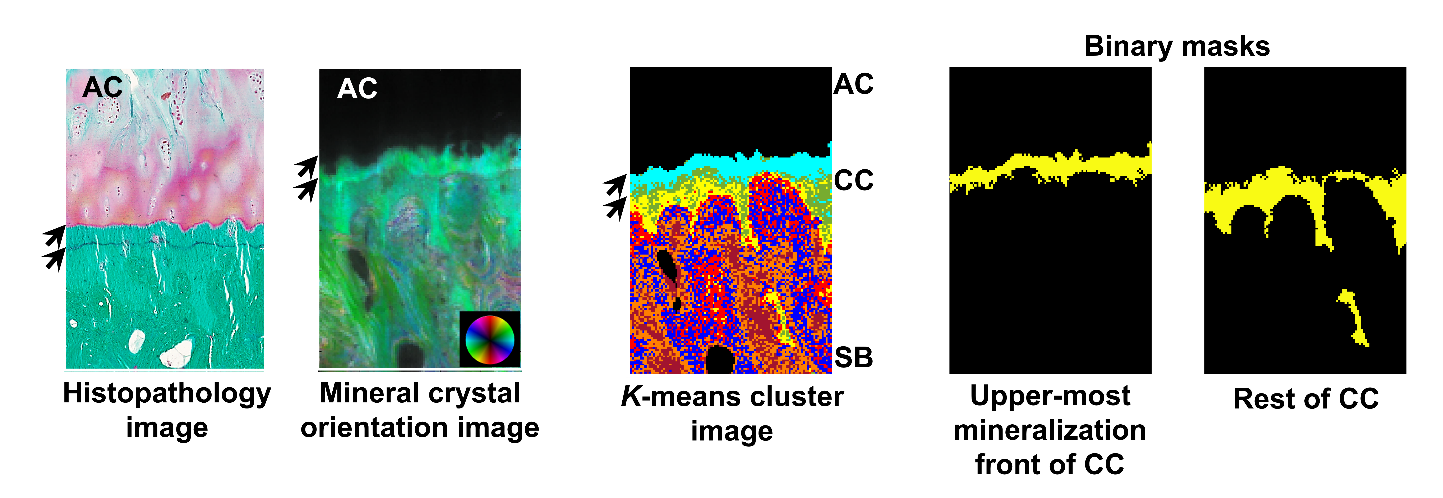
